# CytoLight: A Rapid and Versatile Fluorescent-Based Labeling Method for Extracellular Vesicle Characterization and Tracking

**DOI:** 10.64898/2026.02.10.705037

**Authors:** Ido Rosen, Ella Itzhaki, Ayala Gover-Proaktor, Saar Shapira, Shirly Partouche, Lamis Qassim, Shay Grinshpan-Langman, Arwa Qasim, Daniella Levy Erez, Feba John, Ziv Porat, Neta Moskovits, Romy Zemel, Tali Ben Zur, Pia Raanani, Dani Offen, Galit Granot, Aladin Samara

## Abstract

Efficient, aggregation-free extracellular vesicles (EVs) labeling is essential for studying their dynamics *in-vitro* and *in-vivo*. However, traditional dyes introduce limitations including aggregation, membrane intercalation, fluorescence transfer and inconsistent performance across EV sources thus distorting quantification, altering surface properties and confounding uptake and biodistribution analyses. Here, we systematically evaluated CytoLight, a luminal dye traditionally used for live-cell imaging, as an alternative for EV quantification, characterization, uptake analysis and *in-vivo* tracking, benchmarking it against PKH26, CFSE and ExoBrite across multiple platforms. CytoLight generated stable, intravesicular fluorescence without aggregation or membrane alteration, eliminating artifacts characteristic of conventional dyes. Using fluorescence-NTA and single-EV flow cytometry, CytoLight showed more consistent labeling across EV types than CFSE or ExoBrite, while avoiding PKH-related micelle-driven artifacts and exhibited compatibility with CD81 dual-detection. In uptake assays, CytoLight produced EV-specific endocytosis-dependent internalization signals exceeding labeled-BPS/protein controls. *In-vivo*, CytoLight-labeled EVs enabled fluorescent biodistribution mapping showing conventional EV tropism patterns distinguishable from labeled-PBS controls. These findings establish CytoLight as an effective, aggregation-free EV-labeling strategy. Its stability, specificity, compatibility with single-EV platforms and reliable performance in both cellular uptake and biodistribution studies position CytoLight as a practical, scalable alternative to current dyes, providing a stronger foundation for standardized and reproducible EV research.

## 1. Introduction

Extracellular vesicles (EVs) are nanosized membrane-bound particles secreted by all living cells, involved in intercellular communication. Their unique properties such as biocompatibility, targeting capability, lack of proliferation capacity, low immunogenicity and low toxicity, combined with the potential to modify their surface and cargo, make EVs valuable for both diagnostic and therapeutic applications. In addition, their presence in various biofluids provides insight into cellular states under physiological and pathological conditions, supporting their use as biomarkers for disease detection^1–3^.

Despite their potential, the widespread clinical application of EVs is currently hindered by several technical and biological challenges, particularly their isolation, characterization and quantification. Current isolation techniques struggle to remove contaminants, such as protein complexes and lipoproteins, which can affect the purity and functionality of EV preparations. Furthermore, the pronounced heterogeneity of EV subpopulations complicates their precise quantification and tracking, necessitating the development of advanced methodologies for reliable EV analyses. Addressing these limitations is critical for enhancing the clinical translation of EV-based diagnostics and therapeutics^4–6^.

Accurate EV labeling is imperative for characterizing EV subpopulations, for elucidating uptake mechanisms and for tracking biodistribution, yet methodological constraints persist. Various approaches are currently employed for EV labeling and tracking, including bioluminescence, nuclear imaging and tomographic techniques, each offering distinct advantages in sensitivity and spatial resolution, but often requiring complex instrumentation and radioisotope handling^7–10^. The most common approach for characterization and tracking of EVs is fluorescence-based labeling, which enables the visualization and quantification of EVs with minimal interference to their physicochemical and biological properties^11–13^. Fluorescently labeled EVs facilitate real-time monitoring of cargo loading, uptake by recipient cells and biodistribution in biological systems, providing valuable insights into EV functionality. Fluorescent labeling may be achieved either through genetic engineering, by fusing EV-specific transmembrane proteins (e.g., tetraspanins like CD63 and CD81) to fluorescent reporters like GFP^14^, or by chemical methods using lipophilic-membrane and -cytoplasmic dyes. While genetically introduced reporters offer high specificity and subpopulation resolution, their implementation demands genetic manipulation and is inherently constrained by specific marker expression patterns.

Fluorescent membrane labeling using lipophilic dyes like PKH26, DiI, DiO, ExoBrite and CellMask represents a widely utilized approach for labeling EVs. Such dyes readily integrate into the phospholipid bilayer of EV membranes, thereby enabling their detection and tracking *via* microscopy or flow cytometry^15–17^. While widely used due to their strong signals and ease of application, membrane dyes may potentially fall short of the “minimal interference” ideal. Non-specific binding, dye aggregation and potential alterations of EV surface properties can affect biological interactions, uptake efficiency and targeting capabilities^18^. Another concern is the tendency of some membrane dyes to self-aggregate, forming contaminating particles with sizes similar to EVs, thus distorting size distribution measurements and compromising quantification reliability^19,20^. Furthermore, many membrane dyes exhibit fluorescence transfer between labeled EVs and surrounding membranes, complicating uptake and tracking studies^21^. Since membrane dyes exclusively stain the lipid bilayer, they provide limited insights into EV cargo distribution and dynamics, making them less useful for functional studies.

Cytoplasmic dyes like carboxyfluorescein succinimidyl ester (CFSE), on the other hand, offer a more reliable alternative for EV labeling while preserving vesicular integrity. Unlike membrane dyes, cytoplasmic dyes penetrate the EV membrane and label the intravesicular space, enabling more accurate assessment of EV cargo distribution and transfer efficiency upon uptake by recipient cells.

CytoLight Rapid Red is a fluorescent dye designed to selectively stain the cytoplasm of live cells without affecting membrane integrity^22–25^. CytoLight provides stable fluorescence *via* a rapid process and is compatible with various imaging techniques. Unlike traditional membrane dyes, it minimizes aggregation artifacts and prevents dye transfer between cells making it a powerful and versatile candidate for EV labeling for quantification, characterization and tracking studies. Given these advantages, we hypothesized that CytoLight’s properties would make it a robust and versatile EV-labeling dye capable of minimizing artifacts associated with existing dyes. To test this, we performed a systematic comparison of CytoLight with three widely used EV dyes, PKH26, CFSE and ExoBrite, assessing their performance in terms of specificity, labeling efficiency and practical applicability across key analytical applications. By addressing persistent challenges in EV labeling and quantification, this study aims to establish CytoLight as a reliable dye for advanced standardized, aggregation-free fluorescence-based EV analysis, supporting both basic and translational EV research.

## 2. Materials and Methods

### 2.1. Cell culture

NK92MI and K562 cell lines were obtained from ATCC (USA). HEK293FT (293) cells were obtained from Invitrogen (USA). C6 cells were a kind gift from Prof. Joseph Kost, Ben-Gurion University. 293, VSVG-GFP-293 (prepared with the VSVG-GFP-puro plasmid from VectorBuilder), C6 and K562 cell lines were maintained in high-glucose Dulbecco’s modified eagle medium (DMEM; Gibco, USA) supplemented with 10% fetal bovine serum (FBS; Biowest, France) and 1% penicillin–streptomycin (Gibco). NK92MI cells were cultured in alpha-minimum essential medium (MEM), supplemented with 12.5% FBS, 12.5% donor horse serum, 0.2mM myo-inositol, 2% OTC human AB serum, 0.1mM 2-mercaptoethanol and 0.02mM folic acid. Cells were cultured at 37°C in a humidified 5% CO□ atmosphere and routinely confirmed to be mycoplasma-free.

### 2.2. EV isolation

NK92MI, 293 and VSVG-GFP-293 cells were cultured in their respective growth media supplemented with the relevant EV-depleted sera. Sera were depleted of endogenous EVs by ultracentrifugation (UC) at 110,000×g for 18h using a Ti-45 rotor (Beckman Coulter, USA) at 4°C, followed by filtration through a 0.22μm PVDF membrane (Merck, Germany). Conditioned media were collected 48-72h after the last medium change. 293-derived EVs (293EVs) and VSVG-GFP-293-derived EVs (GFPEVs) were isolated by an UC-based protocol. Briefly, media were sequentially centrifuged at 200×g for 5 min, 900×g for 10 min and 10,000×g for 30 min. The supernatant was filtered through a 0.22μm PVDF membrane and ultracentrifuged at 110,000×g for 4h using a Ti-45 rotor. For NK92MI-derived EVs (NKEVs), conditioned medium was filtered through a 0.22μm PVDF membrane and subsequently concentrated approximately 30-fold by tangential flow filtration (TFF) using a 300kDa hollow-fiber membrane (SP-D04-E300-05-N; Repligen, USA) with a constant transmembrane pressure of 0.3 bar, followed by 5x diafiltration with PBS filtered with a 0.1μm PVDF membrane (Merck), reconcentrated up to 40-fold the original volume and subsequently loaded onto an Izon qEV10/35nm size exclusion chromatography (SEC) column, (Izon Science, New Zealand) equilibrated with 0.22μm filtered PBS. Three 5ml fractions containing the EV-enriched eluate were collected and pooled. EV preparations were immediately stored at −80°C and thawed up to 48h before use and kept refrigerated at 4°C.

### 2.3. Fluorescent labeling of EVs

#### 2.3.1. CytoLight

Unless otherwise stated, EV preparations were resuspended in 50μl particle-free PBS (0.22μm filtered) and labeled with CytoLight at a final concentration of 1-10μM (0.5-5μl of 100μM stock), followed by incubation for 30 min at 37°C. For NTA characterization, labeled EVs were diluted in particle-free PBS to a residual free-dye concentration of 0.005–0.01μM prior to measurement. For experiments requiring removal of unbound dye, samples were subjected to UF using Amicon Ultra 30kDa MWCO regenerated cellulose filters (Merck) and centrifuged at 14,000×g for 10 min at 4°C, after which the retentate was washed once with 0.22μm filtered PBS under the same conditions and recovered for downstream use. All applied controls were processed in parallel and included dye-only controls (CytoLight in particle-free PBS subjected to the same UF or dilution steps), protein-specificity controls (EV-depleted serum matched to the protein concentration of the EV sample and labeled with CytoLight) and detergent lysis controls (CytoLight-labeled EVs incubated with 2% NP-40 for 30 min to disrupt membranous particles).

#### 2.3.2. PKH26

PKH26 labeling was performed according to the PKH26 Red Fluorescent Cell Linker Kit (Merck) manufacturer’s protocol. Briefly, 100µl EVs were diluted in 900µl Diluent C and 6µl PKH26 dye were subsequently added. The mixture was pipetted for 30 sec and then left to rest for 5 min at room temperature. Free dye was quenched by adding 2ml of 10% BSA in 0.22μm filtered PBS. The volume was brought to 8.5ml with serum-free DMEM. Sucrose solution (1.5ml of 0.97M) was pipetted slowly into the bottom of the tube. The tubes were centrifuged at 190,000xg for 2h at 4°C. The media and interface layers were aspirated, and the pellet was resuspended in 0.22μm filtered PBS and transferred to an Amicon ultra 10kDa MWCO regenerated cellulose filters (Merck). Thereafter, 9ml filtered PBS and 0.75ml media were added and the Amicon was spun at 3,000xg until volume was reduced to 0.5ml.

CFSE: EVs were stained with 20µM CFSE and incubated for 2h at 37°C. Free dye was removed either by UC at 190,000×g for 2h at 4°C using an SW 41 Ti rotor, or ultrafiltration (UF) using Amicon ultra 30kDa MWCO regenerated cellulose filters (Merck) at 14,000×g at 4°C for 10 min, followed by one wash with 0.22μm filtered PBS.

#### 2.3.3. ExoBrite

One vial of ExoBrite 640/660 CTB EV staining kit (Biotium, USA) component A was diluted in 100µl ExoBrite reconstitution solution. This solution was aliquoted and kept refrigerated at 4°C. EVs were labeled according to the manufacturer’s recommendations.

### 2.4. Nanoparticle tracking analysis (NTA)

Particle size distribution and concentration were determined using a ZetaView X30 TWIN instrument (Particle Metrix, Germany). The ZetaView system was calibrated with polystyrene particles with a known average size of 100nm (Particle Metrix). Automated quality control measurements were performed prior to sample measurements. Prior to measurement, EV samples were equilibrated at room temperature and diluted to the optimal concentration for measurement in filtered PBS.

Standard scatter mode measurements (sc-NTA) used factory default settings with a 488nm laser. All samples were examined under identical settings of temperature between 22-25°C, sensitivity 75, shutter speed 100 and frame rate of 30 frames/sec.

Single wavelength fluorescence NTA (fl-NTA) employed a 488nm laser with a 500nm long-pass filter for CFSE or PKH26 detection, and a 640nm laser with a 660nm long-pass filter for CytoLight or ExoBrite detection. Samples were examined under identical settings of temperature between 22-25°C, shutter speed 100 and frame rate of 30 frames/sec. Sensitivity for PKH26, CFSE and ExoBrite measurements was set at 95 and for CytoLight measurements was set between 83-95.

For colocalization measurements, GFPEVs were labeled with CytoLight as described above, creating dual-labeled EVs (GFP, 500nm filter; CytoLight, 660nm filter). The negative control consisted of a mixture of two separate EV populations with comparable event numbers: a GFP-positive and a CytoLight-positive population. Fluorescent beads (0.2µM PS 640/660nm; Particle Metrix) were used as a positive control. To certify that the fluorescent-positive events are indeed vesicles, designated purified EV samples were incubated with 2% NP-40 for 30 min.

For each measurement, three replicates were performed by scanning 11 positions and removing outlier positions. Acquisition and analyses were performed by ZetaView software (version 8.05.16 SP7, Particle Metrix) for all single-wavelength experiments. For colocalization experiments, ZetaNavigator (version 1.4.7.6, Particle Metrix) was used for acquisition, and ParticleExplorer (version 4.3.4.4, Particle Metrix) was used for analysis.

### 2.5. EV immunostaining and single-EV flow cytometry (EV-FCM)

For surface phenotyping, 2μl of anti-CD81 antibody (anti-human, Vio® Bright B515, REAfinity, Miltenyi Biotec, Germany) or isotype control (REA control antibody, human IgG1, Vio® Bright B515, Miltenyi Biotec) were added to 10μl of PBS with or without EVs and incubated overnight at 4°C in the dark. EV-FCM was performed using either CytoFLEX nano flow cytometer (Beckman Coulter) or ImageStreamX Mk II imaging flow cytometer (Cytek Biosciences, USA). Side-scatter triggering and fluorescence detection parameters were optimized using the manufacturer’s default quality control protocol along with their proprietary scatter spheres (C85323, Beckman Coulter) and fluorescent spheres (C85324, Beckman Coulter). To certify that the fluorescent-positive events are indeed vesicles, designated purified EV samples were incubated with 2% NP-40 for 30 min. Data of CytoFLEX nano and ImageStreamX Mk II were analyzed with CytExpert nano software (version 1.1.0.6, Beckman Coulter) and IDEAS software (version 6.3, Cytek Biosciences), respectively.

### 2.6. Immunocytochemistry and confocal microscopy

C6 cells were seeded on glass coverslips at 30,000 cells/well in 24-well plates, cultured for 24h and then incubated for 12h in the dark with either 10^9^ CytoLight-labeled NKEVs, 10^9^ unstained NKEVs or CytoLight-labeled PBS at a final volume of 500μl. Cells were washed with PBS, fixed in 4% paraformaldehyde, permeabilized with 0.1% Triton X-100 and blocked with 5% BSA. Immunostaining was performed overnight at 4°C using rabbit anti-EAAT1 (ab416, Abcam, UK) used as a membranal marker and mouse anti-TSG101 (clone 4A10, Thermo Fisher Scientific, USA) used as an endosomal marker, each at 1:500 dilution in 5% BSA blocking solution overnight. Alexa Fluor-conjugated secondary antibodies (488 anti-rabbit A-11008 and 561 anti-mouse A-11018, respectively; Invitrogen) were applied at 1:1,000 dilution for 1h at room temperature. Nuclei were counterstained with DAPI (1:1,000; Merck) for 5 min. Slides were imaged using a confocal microscope (Eclipse Ni-E, Nikon Corp., Japan), using a 20x NA 1.4 objective or 100x NA 0.75 objective, with NIS-Elements AR (Version 5.40.02, Build 1659, 64-bit; Nikon corp.). Images were analyzed using ImageJ (Version 1.54p; National Institutes of Health, USA). CytoLight-TSG101 colocalization analysis was performed using the JacoP plugin (version 2.1.4) to calculate Costes’ randomization based colocalization^26^.

### 2.7. Flow cytometry

C6 and K562 cells (10^4^ and 6×10^4^ cell/well, respectively) were seeded in 96-well plates and incubated with 10^9^ labeled EVs in a final volume of 200μl for 12h in the dark. Unstained EVs and labeled PBS were applied as controls. PBS containing EV-depleted serum with a protein concentration equivalent to that measured in the EV sample was applied as a control for CytoLight’s background staining. Alternatively, C6 and K562 cells were pre-treated with 80μM Dynasore (Merck) or with vehicle (0.1% DMSO) for 30 min and subsequently incubated with labeled EVs and controls as described above. Following the incubation, cells were collected, washed twice with PBS, resuspended in PBS and analyzed by flow cytometry (Gallios, Beckman Coulter). All samples were read under identical hardware settings and analyzed by Kaluza Analysis software (version 2.4, Beckman Coulter).

### 2.8. In-vivo biodistribution studies

Male non-obese diabetic (NOD)-Rag1null IL2rγnull (NRG) mice (10 weeks of age) were administered 10^11^ CytoLight-labeled NKEVs or volume-matched CytoLight-labeled PBS *via* tail vein injection. Four and twenty-four hours post injection, animals were euthanized and liver, spleen, kidney, colon, lung, brain, heart and blood were harvested, rinsed in PBS and imaged *ex-vivo* using an IVIS Lumina III imaging system (PerkinElmer, USA) with the fluorescence settings Fstop = 2, binning = medium, exposure = auto, lamp level = high. Liver tissues were also mechanically dissociated to generate single-cell suspensions for flow cytometric analysis of CytoLight fluorescence. All experimental procedures were carried out in accordance with the protocol approved by Institutional Animal Care and Use Committee (RMC-IL-2501-105-5).

### 2.9. Statistical analysis

Quantitative data are presented as mean ± standard error from at least three independent biological replicates. Unpaired two-tailed Student’s t-tests were applied in comparisons between two groups, assuming similar variance when deemed appropriate by F-test, and unequal variance otherwise. Multiple group comparisons, performed on R (R foundation for statistical computing), employed Bartlett’s test to ensure similar variances, then one-way ANOVA, followed by Tukey’s post-hoc test. Statistical significance cutoff was defined as p≤0.05. Histograms and scatter-plots were generated by Kaluza analysis software (version 2.4). CytoFlex nano plots in the Suppl. file were generated by CytExpert nano (version 1.1.0.6; Beckman-Coulter). Animal study IVIS box plots were made with R (R foundation for statistical computing). All other plots were generated using Excel (Microsoft, USA). Hill-function curve fitting was performed in Python (PSF, USA) using the SciPy curve fit function to model the fluorescence-to-scatter ratio as a sigmoidal response according to the Hill equation (y = Emax × [x^n^/(EC50^n^ + x^n^)])^27^.

## 3. Results

### 3.1. CytoLight effectively labels EVs

Our initial experiments aimed to assess the feasibility of CytoLight to effectively label EVs. To establish an optimal staining window for accurate EV quantification by fluorescence NTA, CytoLight-labeled EVs were analyzed in parallel by scatter-mode NTA (sc-NTA), reporting the total particle concentration and fluorescence-mode NTA (fl-NTA), detecting only dye-positive particles. To identify the optimal conditions for CytoLight EV labeling, we sought to balance three key parameters: achieving a sufficiently strong fluorescent signal, minimizing background from unbound dye and maintaining adequate EV concentration for reliable detection. To this end, NKEVs, isolated by TFF followed by SEC (TFF-SEC), were first incubated with increasing CytoLight concentrations (1, 5, and 10µM) for 30 min at 37°C, to allow dye uptake followed by serial dilution to yield measurement optimal concentrations spanning 0.005–1µM, to proportionally reduce free dye while preserving the signal from labeled vesicles. Particle free PBS containing equivalent serially diluted dye concentrations was used to prepare a titration curve.

NTA analysis revealed that CytoLight labels EVs in a quantifiable manner when low concentrations of free-dye are present (≤0.01µM), where a clear fraction of fluorescent-positive particles was detected at a ratio of approximately 0.3–0.7 fluorescent particles to total particles (Fig. 1A, left axis). At higher free dye concentrations (≥0.025µM), we observed fluorescence-to-particle ratio that vastly exceeded 1.0 (up to 3.2□±□14.9), indicating substantial interference from background signal due to unbound dye. Beyond this threshold, particularly above 0.05µM, discrete fluorescent particles could no longer be resolved (Fig. 1A, left axis), probably due to pronounced signal saturation from excess free CytoLight molecules in the solution (Fig. 1A, right axis). An exception was observed in samples initially labeled with 10µM CytoLight, which still exhibited quantifiable fluorescent particles. This suggests that higher labeling concentrations enhance dye incorporation into vesicles, yielding fluorescent signals that remain distinguishable from background noise. These findings were in line with the fluorescent signal scattering profile of dye-only PBS samples, which exhibited marked optical interference at equivalent CytoLight concentrations (Fig. 1A, right axis). PBS samples containing 0.025-1µM CytoLight showed high fluorescent scattering intensities (74 ± 7.3 to 255), indicative of strong optical saturation, whereas at 0.005-0.01µM, displayed markedly lower intensities (7.6 ± 0.8 to 34 ± 6), defining the range in which the background signal no longer dominated the measurement window.

**Fig. 1.**
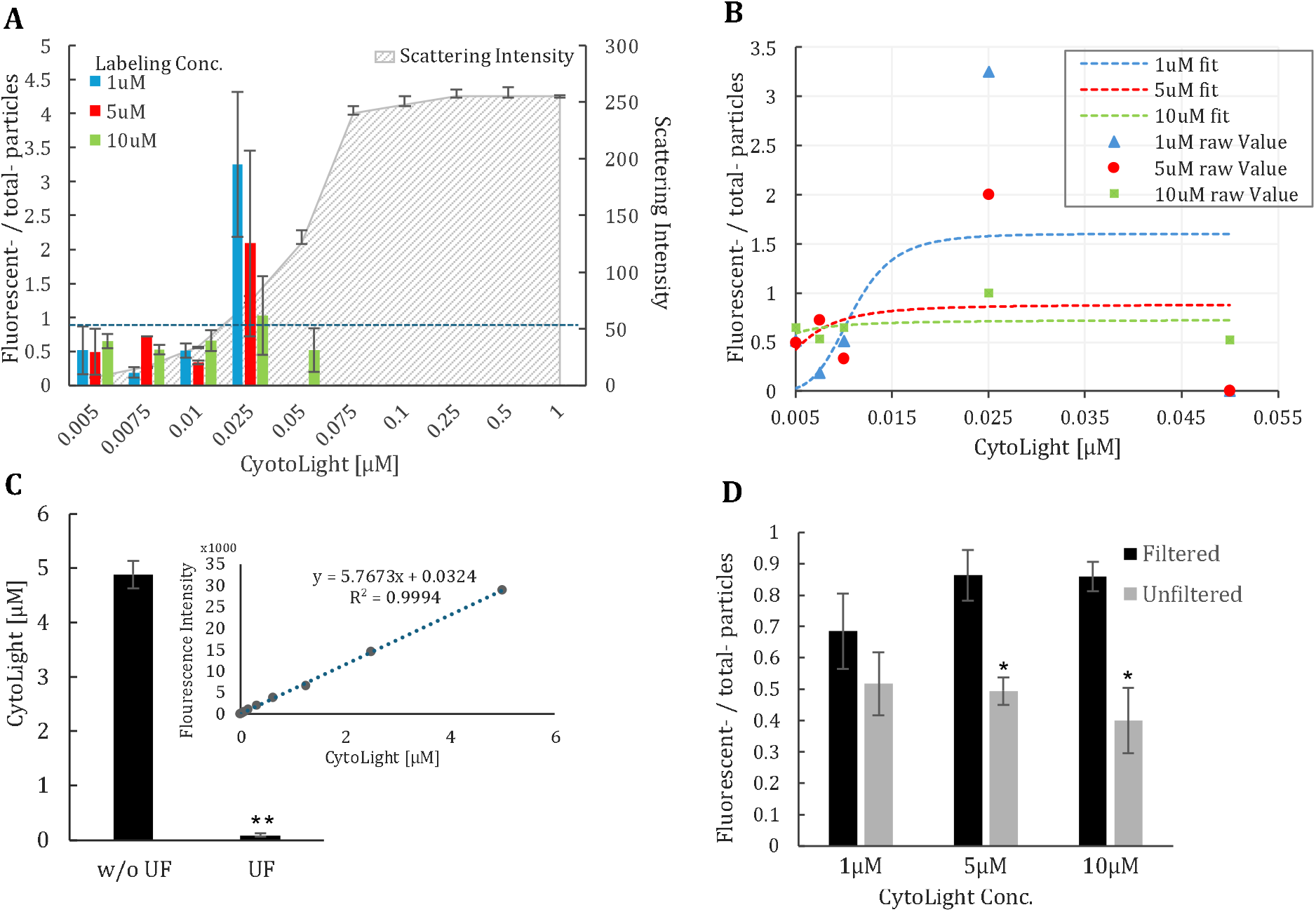
Optimization of CytoLight labeling concentration and removal of excess dye. NKEVs were incubated with 1, 5 and 10µM CytoLight for 30 min at 37□. Subsequently each sample was diluted to an array of concentrations ranging from 0.005-1µM and analyzed simultaneously using fl-NTA and sc-NTA. Unstained NKEVs and CytoLight labeled PBS were applied as controls. **(A)** Scattering intensity of PBS samples (gray shading, right axis) and ratio of CytoLight-positive to total particles in stained EV samples (bars, left axis) across increasing dye concentrations. The X axis represents CytoLight concentrations after dilution. **(B)** Hill-function for ratio values for 1µM (blue), 5µM (red) and 10µM (green) labeling arrays. Experimental points are plotted with fitted curves (dashed lines). **(C)** PBS was labeled with 5µM CytoLight and measured before and after UF. Fluorescence intensity was quantified using a microplate reader. Shown is a bar graph of effective dye concentration before and after UF. CytoLight concentrations were determined using a calibration plot of fluorescence *vs*. dye concentration (inset). **(D)** Ratio of fluorescent to total EVs in filtered *vs*. unfiltered samples for 1, 5 and 10µM labeling reactions. Values represent a mean of at least 3 experiments ± SEM. * P≤0.01, ** P≤0.001.

Hill-function fitting of the fluorescence-to-scatter ratios revealed distinct kinetic profiles for the three labeling conditions (Fig. 1B). For the 1µM labeling arm, the fluorescence response displayed pronounced cooperativity (E□□□ = 1.62; EC□□ = 0.0109µM; Hill slope = 5.0), reflecting a signal-limited regime characterized by a sharp transition in detectable labeling. The 5µM arm yielded a broader, more gradual curve (E□□□ = 0.89; EC□□ = 0.0053µM; Hill slope = 2.37), consistent with a background-limited regime, in which free dye contributes optically to total fluorescence. In contrast, the 10µM arm reached lower maximal ratios (E□□□ = 0.72; EC□□ = 0.0018µM; Hill slope = 1.51), defining a saturation-stable regime dominated by dye-loaded vesicles that maintain stable brightness across a broad dilution range. Importantly, 10µM-labeled samples remained quantifiable even when only diluted to 0.05µM CytoLight, supporting the notion that increased dye loading per vesicle improves detectability despite background fluorescence.

At higher CytoLight concentrations, apparent labeling efficiency decreased due to strong optical interference from free dye, which inflated fluorescence readings, generated false-positive particle detection, and ultimately rendered measurements unreliable above 0.025µM free dye concentration (Fig. 1A, B), posing a challenge for downstream applications, such as EV uptake assays and *in-vivo* studies which require highly fluorescent EVs to achieve detectable signals. To address this limitation, we established and optimized a UF-based purification step for the removal of excess, unbound dye from the samples. PBS samples containing 5µM CytoLight were subjected to UF through regenerated cellulose membranes (30kDa cutoff). As shown in Fig. 1C, UF removed 98.16 ± 0.7% of free dye. Notably, when applied to EV-containing samples, UF markedly reduced optical interference caused by nonspecific fluorescence (as illustrated in Fig. 1A, right axis) enabling more accurate readings of true EV events (Fig. 1D) while preserving total scatter counts and modal particle size (shown in the comparative analysis provided in Fig. 3B), indicating that CytoLight–EV complexes remain stable throughout the purification process. These results confirm that UF efficiently removes free dye and minimizes optical background, offering a rapid and reproducible alternative to time-intensive UC-based cleanup methods^28^, which have also been reported to induce EV aggregation and significant particle loss^29,30^.

Taken together, these results demonstrate that CytoLight effectively labels EVs when applied at micromolar concentrations (1–10µM), and that optimal and quantifiable fluorescence is achieved at diluted free dye concentrations of 0.005–0.01µM, eliminating the need for additional purification. For background-sensitive downstream applications, however, signal interference can be effectively mitigated by a brief UF step, which restores measurement accuracy without compromising EV integrity.

Having established that CytoLight effectively labels EVs, we next compared its performance with the commonly used EV dyes PKH26, CFSE and ExoBrite, focusing on two critical parameters: (i) their tendency to form aggregates; and (ii) their efficacy in labeling EVs.

### 3.2. Aggregation propensity of CytoLight

The formation of dye aggregation is a common artifact of EV dyes that generates false-positive fluorescent particles mimicking stained EVs in NTA analysis. To compare CytoLight’s tendency to form aggregates to that of commonly used EV dyes, PBS was stained with each of the dyes using concentrations equivalent to those typically used for labeling 10^9^ EVs. As seen in Fig. 2A, PBS stained with PKH26 consistently contained a high number of PKH26-labeled nanoparticles (1.25×10^1^□ particles/ml) with ∼58% of these particles exhibiting fluorescence in fl-NTA (Fig. 2A, B). sc-NTA revealed a particle size distribution ranging from 70-400nm, resembling that of EVs (Fig. 2C). In contrast, PBS labeled with CytoLight, CFSE or ExoBrite showed only trace amounts of non-fluorescent and fluorescent particles, indicating minimal to no self-aggregation for these dyes (Fig. 2A, B). These results were further confirmed by EV-FCM analysis, where CytoLight-, CFSE- and ExoBrite-labeled PBS showed minimal amounts of fluorescent particles. In contrast, PKH-labeled PBS showed nearly 75% fluorescent particles and displayed a polydisperse particle size distribution resembling that of EVs, underscoring the risk of mistaking dye aggregates for genuine vesicles and thereby compromising accurate quantification (Fig. 2D). These findings highlight the inherent limitation of PKH26, which generates dye-derived nanoparticles. Consequently, PKH26 was excluded from further analyses.

**Fig. 2.**
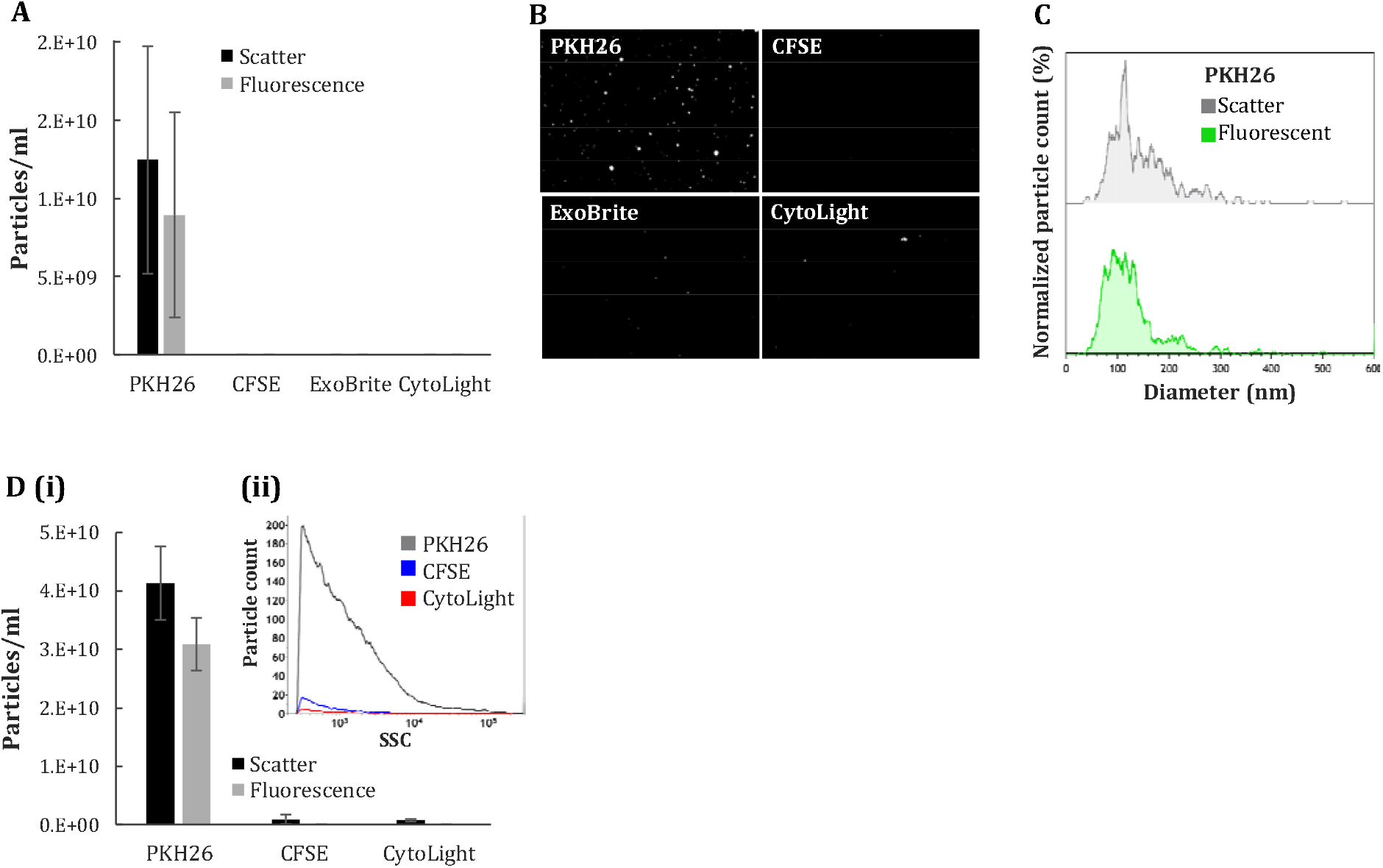
CytoLight presents with minimal to no aggregation. PBS was labeled with PKH26, CFSE, ExoBrite and CytoLight and analyzed according to their respective protocols. Self-aggregation was assessed by NTA and EV-FCM. **(A)** Bar graph of particle concentration as per sc- and fl-NTA. **(B)** Representative images from NTA videos of indicated samples. **(C)** Representative size distribution profiles overlay of PKH26 particles. **(D(i))** Bar graphs of particle concentrations as per EV-FCM. **(D (ii))** Representative size histogram of fluorescent particles. Values represent a mean of at least 3 experiments ±

### 3.3. Labeling efficiency of CytoLight

We next sought to compare CytoLight’s labeling efficiency with CFSE and ExoBrite. To reflect common laboratory workflows, we selected two representative EV models. NKEVs prepared by TFF-SEC and 293EVs prepared by UC. Labeling was conducted in parallel for each dye in accordance with its validated protocol and analyzed by NTA. At 10µM, CytoLight achieved robust labeling in both EV types, achieving high efficiencies of 98% ± 11% CytoLight-positive particles in 293EVs (Fig. 3A, B(i)) and 81% ± 29% in NKEVs (Fig. 3A, B(ii)), without significant alteration in EV size distribution in sc-NTA.

**Fig. 3.**
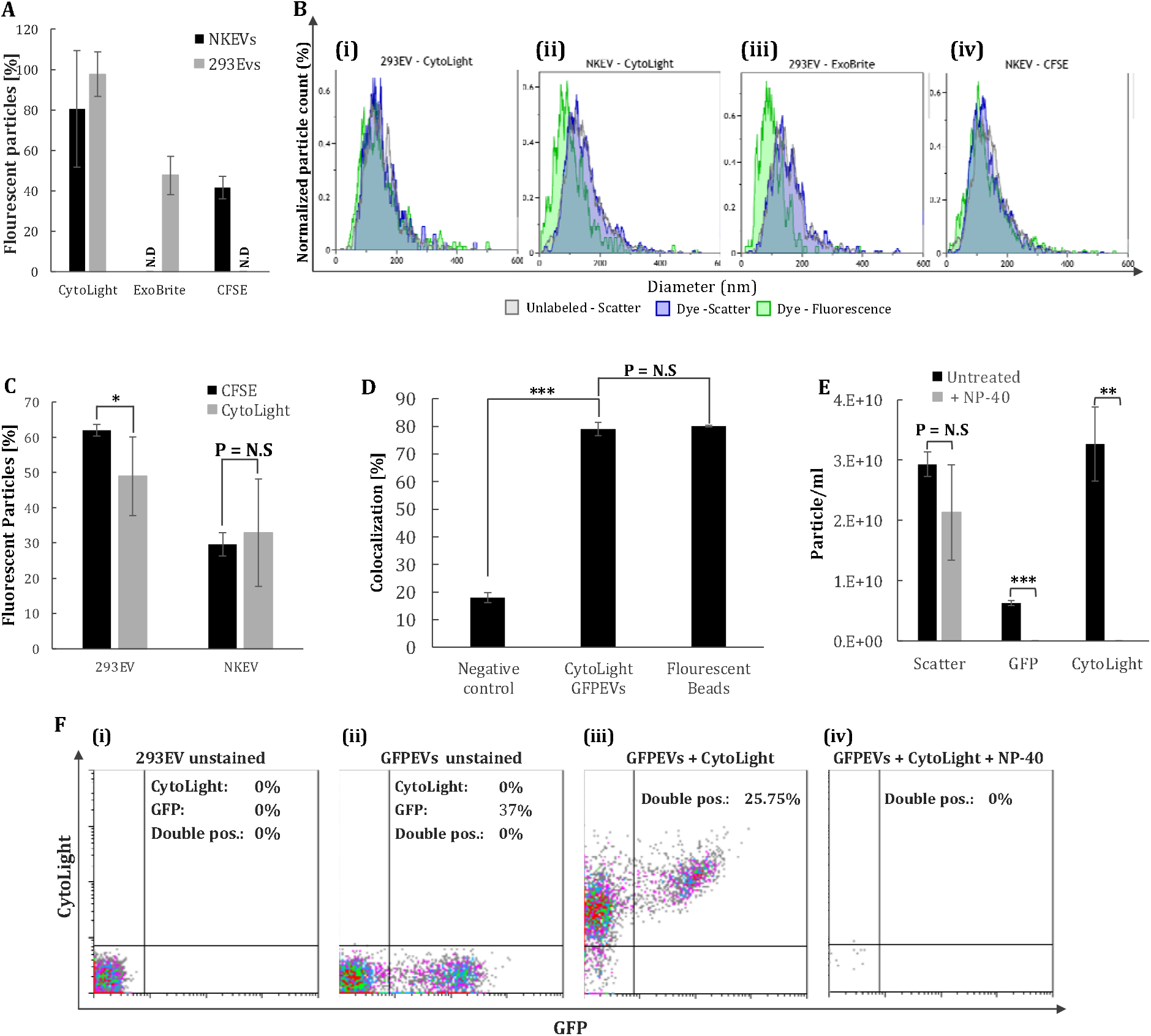
Comparative EV labeling efficacy of the different dyes. 293EVs and NKEVs were labeled with CytoLight, ExoBrite and CFSE, following respective protocols. **(A)** Percentage of fluorescent 293EV and NKEVs quantified by fl-NTA. **(B)** Representative size distribution overlays of unlabeled- and labeled-EVs in sc-mode (grey and blue) and dye-labeled EVs in fl-mode (green). **(C)** EV-FCM analysis of CFSE- and CytoLight-labeled 293EVs and NKEVs, showing percentages of fluorescent particles. **(D)** Colocalization of CytoLight and GFP signals in the indicated particles. A mixture of CytoLight-labeled 293EVs and non-labeled GFPEVs were applied as a negative control. Fluorescent beads were applied as positive controls. Signals were acquired simultaneously with 488nm and 640nm laser. **(E)** Concentrations of CytoLight-labeled GFPEVs treated or not with NP40, quantified by sc- and fl-NTA. **(F)** Representative EV-FCM dual-parameter plots of GFP and CytoLight signals, after gating for singlets: **(i)** Unstained 293EVs; **(II)** unstained GFPEVs; **(III)** stained GFP-EVs; **(IV)** stained GFPEVs after NP-40 treatment. Values represent a mean of at least 3 experiments ± SEM. * P≤0.05, ** P≤0.01, *** P≤0.0001. N.S: not significant.

ExoBrite achieved 48 ± 9% labeling of 293EVs, but only at concentrations 33-fold higher than the manufacturer’s minimal recommendation (Fig. 3A, B(iii)). In NKEVs, no measurable labeling was detected (Fig. 3A). This observation is consistent with the manufacturer’s disclaimer that ExoBrite’s performance is cell-type dependent, yet the requirement for such high dye amounts rendered it impractical for applications requiring large numbers of labeled EVs, and it was not tested further.

CFSE achieved moderate labeling efficiency of NKEVs with 42 ± 6% fluorescent particles without affecting size distribution (Fig. 3A, B(iv)). In the 293EV sample, we noticed substantial background fluorescence with CFSE, precluding reliable quantification (Fig. 3A). Reducing CFSE concentration did not improve the signal-to-noise ratio (data not shown). Taken together, CytoLight proves more consistent, efficient and practical labeling across both EV sources, thereby outperforming CFSE and ExoBrite.

It is worth noting that in all labeled samples, fl-NTA detected smaller particle populations than sc-NTA of the same labeled samples indicating that while CytoLight, ExoBrite and CFSE did not alter membrane properties, they enhanced detection of smaller EVs by improving signal-background ratio and allowing higher NTA sensitivity (Fig. 3B).

To complement the NTA data, CytoLight and CFSE were further tested by EV-FCM using nanoscale flow cytometer CytoFLEX-nano and ImageStream. When analyzed by CytoFLEX-nano, both dyes generated fluorescent signals in 293EVs and NKEVs (Fig. 3C, Suppl. Fig. 1A, B). In 293EVs, CFSE labeled a significantly higher fraction of EVs compared to CytoLight (62 ± 2% *vs*. 49 ± 3%, p = 0.04). In NKEVs, labeling efficiencies were lower overall and comparable between the two dyes (33 ± 11% *vs*. 30 ± 15%, p = 0.9) (Fig. 3C). CytoLight- and CFSE-labeled 293EVs were also detected by imaging flow cytometry (ImageStream), confirming the CytoFLEX-nano findings (Suppl. Fig. 1C, D). These results highlight the compatibility of CytoLight with different EV sources, isolation techniques and analysis methods.

Interestingly, CytoFLEX-nano analyses revealed distinct labeling patterns: CFSE produced a bimodal distribution, separating the EV population into highly fluorescent and non-fluorescent subsets consistent with esterase-dependent activation and heterogeneous esterase content among EVs^30,31^. CytoLight, on the other hand, produced a uniform fluorescence shift toward higher fluorescence intensity, in line with homogeneous and esterase-independent incorporation (Suppl. Fig. 1A, B). This uniform labeling minimizes the biological variability constraining CFSE and enables a more universal quantification across diverse EV populations.

To confirm that CytoLight specifically labels *bona fide* EVs rather than co-isolated proteins or non-vesicular contaminants, we employed several independent validation approaches. To validate the specificity of CytoLight, we utilized EVs isolated from 293 cells engineered to express GFP-fused vesicular stomatitis virus glycoprotein (VSV-G), a known marker for cell-derived particles^33^. Remarkably, NTA analysis revealed virtually complete colocalization between GFP and CytoLight signals: 79 ± 0.02% of GFP-positive particles were also CytoLight-positive, comparable to the 80 ± 0.02% colocalization benchmark of the positive control (p = 0.3), and significantly higher than the negative control (18 ± 0.003%, p = 0.00004) (Fig. 3D). Second, we demonstrated that the membrane-disrupting detergent, NP-40 abolished both GFP and CytoLight signals in CytoLight-labeled GFPEVs, confirming that both observed signals originated from intact vesicular structures (Fig. 3E). EV-FCM further validated and reinforced these findings, showing 100% colocalization of the GFP-CytoLight signals, and their elimination following NP-40 treatment (Fig. 3F). Finally, dual labeling with anti-CD81-FITC demonstrated that 55% of CytoLight-positive events were also CD81-positive (Suppl. Fig. 1E-H), supporting robust labeling of genuine EVs.

### 3.4. CytoLight is suitable for in-vitro EV uptake analysis

Building upon the robust labeling performance of CytoLight, we next evaluated its utility as a tool for EV tracking through uptake assays using both flow cytometry and confocal microscopy. For flow cytometry-based uptake assessment, 10^9^ NKEVs were labeled with 10µM CytoLight or 20µM CFSE. C6 cells were then incubated with labeled NKEVs for 12h. Excess dye was removed by UF. Unstained EVs served as negative controls, and particle-free PBS labeled with the corresponding dyes was processed in parallel to account for background fluorescence and non-specific signal transfer. Mean fluorescence intensity (MFI) was log2 transformed due to the exponential change in MFI and due to the differences in variance between the different groups. Cells exposed to CFSE-labeled NKEVs showed no significant MFI increase over CFSE-labeled PBS controls (1 ± 0.87 *vs*. 0.65 ± 0.46 log2fc, p = 0.72; Suppl. Fig. 2A). Comparable results were obtained when UC rather than UF was used for dye removal (0.25 ± 0.23 *vs*. 0.33 ± 0.12 log2fc, p = 0.78; Suppl. Fig. 2A), or when the EV dose was increased to 5×10^10^ particles (28.9 ± 28.0 *vs*. 36.9 ± 30.8 log2fc, p = 0.63; Suppl. Fig. 2B). These findings suggest that the apparent CFSE signal largely reflects unbound dye rather than EV uptake, aligning with previous reports of its non-specific transfer and ineffective purification by UC or UF^21,31,34,35^. In contrast, C6 cells incubated with CytoLight-labeled NKEVs exhibited significantly higher fluorescence than those exposed to CytoLight-labeled PBS controls (5.67 ± 0.2 *vs*. 0.28 ± 0.06 log2fc, p = 0.00009; Fig. 4A).

**Fig. 4.**
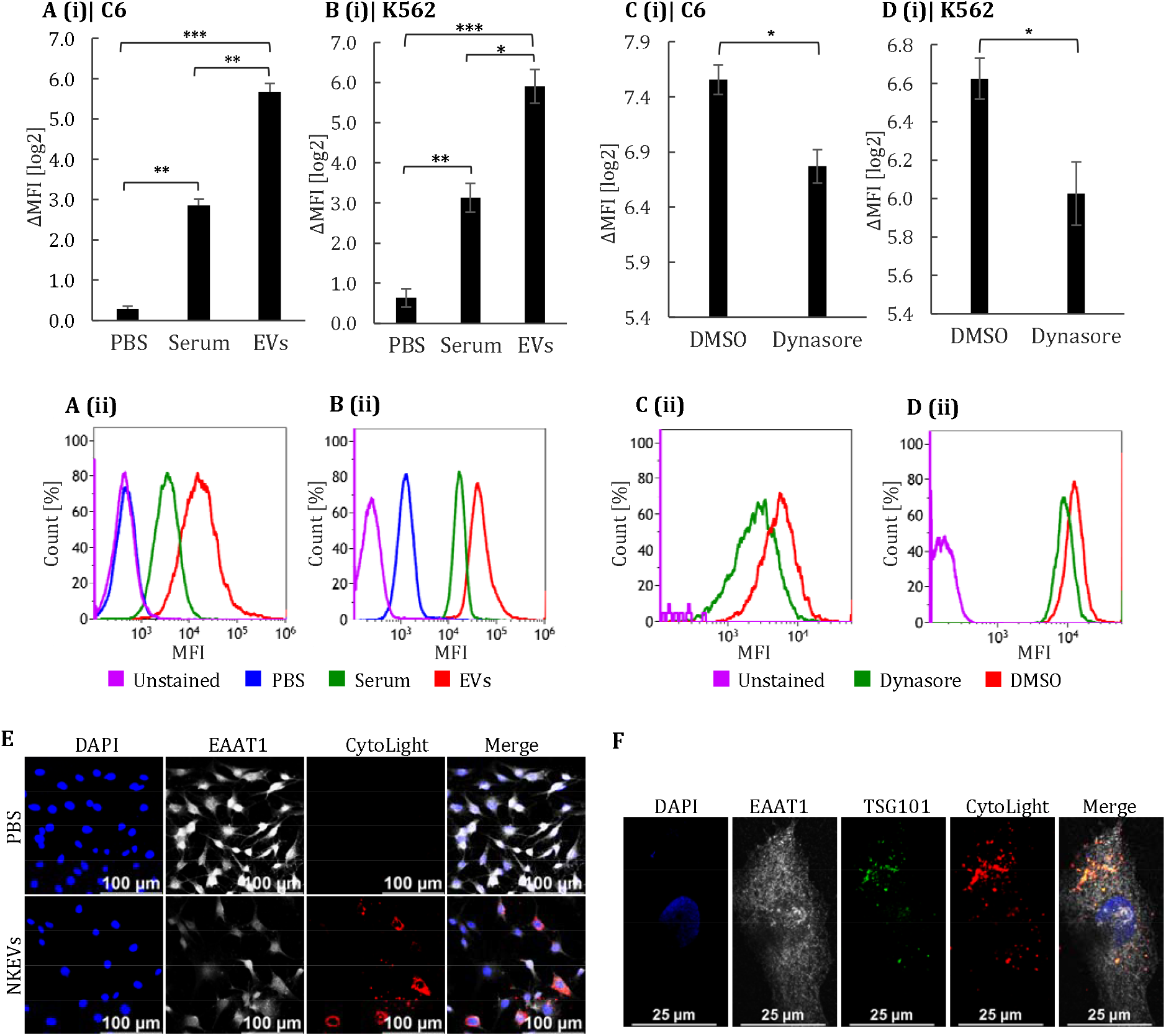
Cellular uptake and intracellular trafficking of CytoLight-labeled EVs. **(A-B)** C6 and K562 cells were incubated with 10^9^ CytoLight-labeled NKEVs for 12h and analyzed by FACS. CytoLight-labeled PBS and serum served as controls for background and nonspecific protein staining, respectively. Shown are bar graphs of log2ΔMFI, relative to control samples treated with unstained EVs **(A(i), B(i))** and representative histograms **(A(ii), B(ii))** of C6 **(A)** and K562 **(B)** cells. **(C-D)** C6 and K562 cells we treated as detailed in A-B and pre-treated with 80µM Dynasore or DMSO. Shown are bar graphs **(C(i), D(i))** and representative histograms **(C(ii), D(ii))** of C6 **(C)** and K562 **(D)** cells. **(E)** C6 cells were incubated with CytoLight-NKEVs or PBS for 12h, and immunostained with the glial plasma membrane marker EAAT1 (grey) and counterstained with DAPI (blue). Representative single-channel and merged confocal images are shown at 20× magnification. **(F)** C6 cells incubated with CytoLight-NKEVs were immunostained with the endosomal marker TSG101 (green) and counterstained with DAPI (blue). Representative single-channel and merged images are shown at 100× magnification. Channels: nuclei (DAPI; blue), membrane (EAAT1; grey), endosome (TSG101; green) and EVs (CytoLight; red). Values represent a mean of at least 3 experiments ± SEM. * P≤0.01, ** P≤0.005, *** P≤0.0001.

One of the chief concerns of EV dyes is non-specific binding to free proteins co-isolated with the EVs. To assess the magnitude of this phenomenon with CytoLight, EV-depleted serum was matched for the protein concentration of the EV sample, and labeled with CytoLight. Although C6 cells incubated with labeled serum were significantly more fluorescent than those incubated with labeled PBS (2.86 ± 0.16 *vs*. 0.29 ± 0.06 log2fc, p = 0.003, Fig. 4A), the signal remained significantly lower than that of labeled-EVs (2.86 ± 0.16 *vs*. 5.67 ± 0.2 log2fc, p = 0.005, Fig. 4A). Similar results were obtained in K562 cells, where CytoLight-labeled EVs induced significantly higher fluorescence compared with both serum (5.9 ± 0.4 *vs*. 3.1 ± 0.35 log2fc, p = 0.02; Fig. 4B) and PBS controls (5.9 ± 0.4 *vs*. 0.63 ± 0.27, p = 0.0005; Fig. 4B). To better understand the mechanism of CytoLight-labeled NKEV uptake, C6 and K562 cells were also incubated with the endocytosis inhibitor Dynasore^36^. Dynasore treatment significantly reduced the fluorescence of cells incubated with labeled-EVs compared to the same cells treated with DMSO in both C6 cells (7.6 ± 0.1 *vs*. 6.8 ± 0.15 log2fc, p = 0.02; Fig 4C) and in K562 cells (6.6 ± 0.2 *vs*. 6.0 ± 0.1 log2fc, p = 0.04; Fig. 4D).

Microscopy-based uptake assessments reinforced these results. In these assays, CytoLight-labeled NKEVs were administered to C6 cells for 12h. Particle-free PBS labeled with CytoLight was used as a control to asses potential background signal. CytoLight-labeled EVs displayed a distinct intracellular punctate pattern of fluorescence in recipient C6 cells, clearly exceeding the signal seen in cells exposed to labeled PBS (Fig. 4E). These punctate colocalized strongly with the endosomal marker, TSG101 (r = 0.7, p<0.0001, Fig 4F), suggesting that CytoLight-labeled EVs were internalized *via* endocytic pathways. These data confirm that CytoLight enables robust and quantifiable EV uptake visualization, with minimal background and biologically relevant patterns.

### 3.5. CytoLight is suitable for in-vivo EV biodistribution studies

CytoLight’s suitability for *in-vivo* EV tracking was next evaluated in NRG mice. NKEVs or PBS controls were labeled with CytoLight, purified by UF and administered intravenously to the tail of healthy male mice. In an initial pilot experiment (n=2), whole-body IVIS imaging revealed fluorescence in multiple organs at 4h and 24h in the mice which received labeled EVs (data not shown). A subsequent study (n=5) confirmed significantly higher fluorescence in liver, colon, kidney, blood, lung and spleen of mice injected with CytoLight-labeled EVs compared to those injected with CytoLight-labeled PBS controls (Fig. 5A, B). In contrast, heart and brain exhibited minimal fluorescence over background. with no significant differences between groups (Fig. 5A, B). Flow cytometry analysis of liver single-cell suspensions further supported these observations, demonstrating markedly increased fluorescence in the CytoLight-labeled EV group relative to the PBS control group (Fig. 5C).

**Fig. 5.**
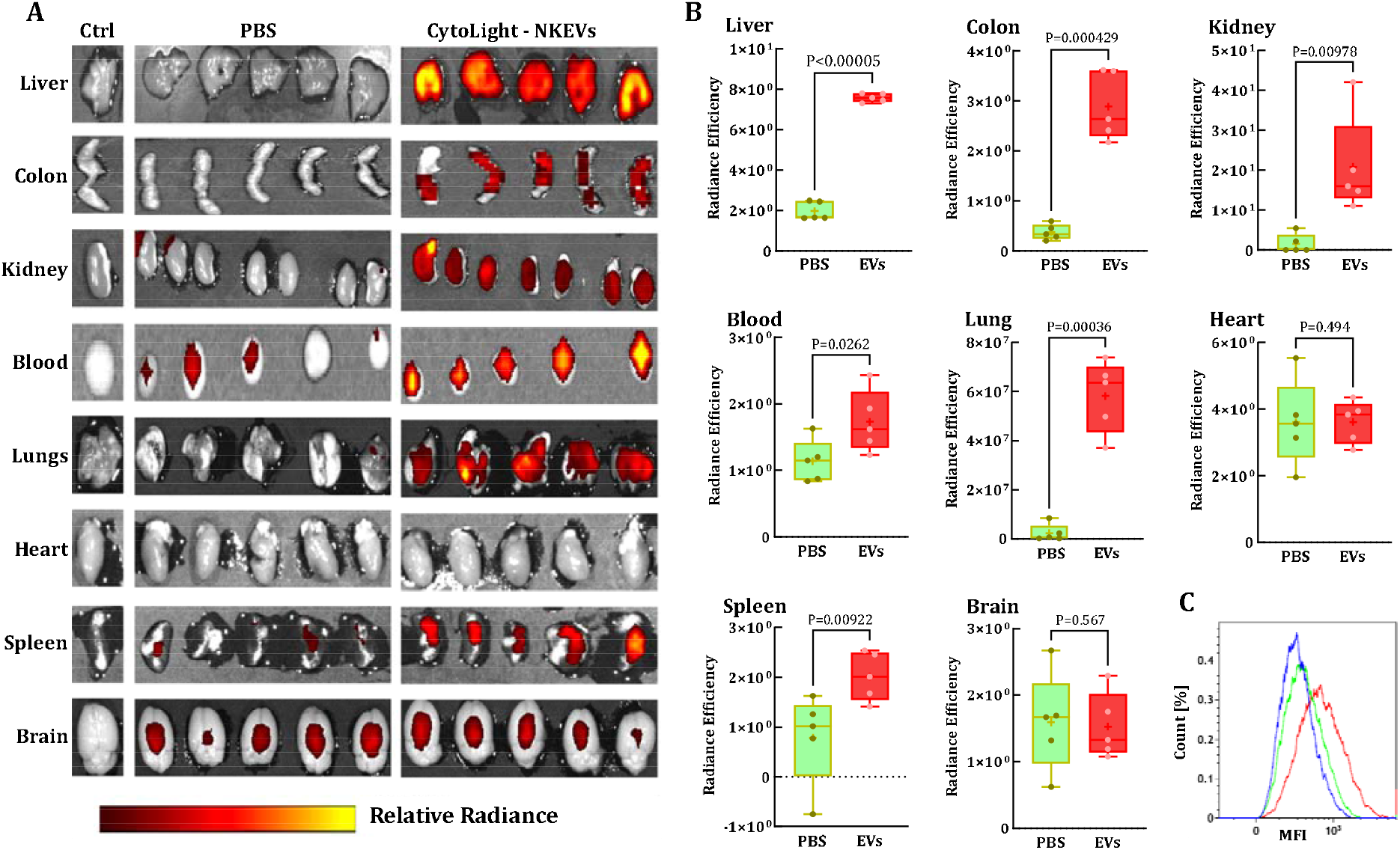
*In-vivo* biodistribution of CytoLight-labeled EVs. NKEVs (10^11^ particles) or particle-free PBS were labeled with CytoLight, purified as previously described and injected intravenously into mice tails (n=5). Organs were collected 24h post-injection and analyzed *ex-vivo* for fluorescence by IVIS. **(A)** Representative *ex-vivo* IVIS images showing relative radiance (relative radiance, [p/s/cm^2^/sr]/[µW/cm^2^]). **(B)** Box plots representing quantification of total radiance efficiency for each organ collected from PBS-injected (green) and NKEV-injected (red) mice. **(C)** Collected livers were dissociated mechanically to single cell solutions and analyzed *via* flow cytometry. Shown are resulting histograms of liver single cell solutions from untreated control (blue), PBS-injected (green) and NKEV-injected (red) mice.

## 4. Discussion

In this study we introduced and systematically evaluated CytoLight Rapid Red as a reliable and versatile fluorescent-based cytoplasmic dye for EV labeling. CytoLight demonstrated robust performance in labeling efficiency, signal specificity and compatibility with both *in-vitro* and *in-vivo* applications across multiple analytical platforms.

The first part of the study pursued evaluating CytoLight’s labeling performance and defining its effective labeling range. Using NTA, we found CytoLight to effectively label EVs when applied at micromolar concentrations (1–10µM), with 5-10µM producing the most stable quantifiable fluorescence signals. However, NTA titration profiles, performed both in PBS and in EV-containing samples, revealed that EVs can only be reliably quantified when residual unbound dye is diluted below 0.01µM as higher levels of free dye dominate the fluorescent readout. Together, these findings defined a two-step operational window at which EVs are best loaded at 5-10µM but must be quantified below 0.01µM residual dye to ensure a strong fluorescent signal that arises from labeled-vesicles rather than free dye. A simple post-labeling dilution step achieved this, preserving intravesicular fluorescence generated during high-concentration labeling while reducing residual dye to within a quantifiable range, thus eliminating the need for subsequent dye removal steps for quantification analyses. Hill-fit analysis of the fluorescence-to-scatter ratios from the titration series further indicated that CytoLight labeling reaches a saturation-limited regime at approximately 10µM, consistent with the plateau observed in the titration curve. This plateau, together with the observed linearity between fluorescence and particle counts (seen at 5µM and more dominantly at 10µM), supports true intravesicular incorporation rather than nonspecific membrane adsorption. Comparable titration profiles have been reported for other dyes (e.g., CFSE, DiO)^37^. As calibration was performed on a single NTA platform, the precise operational range should be re-defined for each instrument.

CytoLight labeling markedly enhanced EV detectability by NTA with fl-NTA in some cases, identifying more CytoLight-labeled particles than could be resolved by standard sc-NTA. This is likely because the dye’s fluorescence increased the signal-to-background ratio, permitting higher sensitivity and improved particle detection. This observation echoes other reports that EV quantification with sc-NTA can underestimate particle concentrations^38,39^. While EV size was unchanged in sc-NTA, confirming unmodified membrane properties, fl-NTA revealed smaller vesicles, suggesting improved sensitivity to dim subpopulations missed by standard analysis. Thus, CytoLight mitigates underestimation biases in conventional sc-NTA, the current standard for EV quantification. However, this observation should be interpreted with caution, as our experiments were conducted without inter-instrument validation^39^.

To assess CytoLight’s performance beyond NTA, we evaluated its behavior on two additional, flow-cytometry based, single-EV detection platforms. On both CytoFLEX Nano and ImageStream, CytoLight-labeled EVs generated fluorescence intensities that were well above instrument background and remained clearly separated from dye-only controls, indicating that the dye provides sufficient photon yield without false-positive and aggregation-driven events characteristic of other commonly used dyes^40^. The fluorescence signal of CytoLight-labeled EVs was abolished after detergent lysis, confirming that the signal originated from intact vesicles rather than free dye or co-isolated protein aggregates. Importantly, CytoLight labeling preserved compatibility with surface phenotyping, enabling co-detection of CD81 on the same EV populations. The convergence of GFP colocalization, NP-40-mediated signal loss, and CD81 co-staining collectively verifies the vesicular specificity of CytoLight signals. These attributes position CytoLight as an ideal reagent for dual-labeling single-EV analysis, facilitating simultaneous intravesicular staining and surface marker profiling in multiparametric cytometry workflows.

Within the above defined operational range and across the quantification platforms used, CytoLight remained stable in suspension, aggregation-free and effectively labeled EVs, without perturbing vesicle size distribution or scattering properties. Once internalized, the dye became intravesicular and impermeant, producing fluorescence confined to individual vesicles with no evidence of dye transfer between particles. This is as opposed to lipophilic dyes, such as PKH26, which intercalate into membranes enabling dye transfer and even promoting vesicle fusion that may alter EV size, surface properties and downstream biological interactions. In addition, PKH26 generates abundant fluorescent aggregates in PBS that are indistinguishable from genuine EVs by size or fluorescence^37,41^. Such aggregates inflate particle counts and distort interpretation of functional downstream analyses^21^. ExoBrite, a cholera-toxin–conjugated membrane-labeling dye, designed to overcome such artifacts, showed inconsistent performance. Labeling was detected in 293EVs only at 33-fold higher concentration than the recommended minimum and was undetectable in NKEVs. This strong cell-type dependence, coupled with high cost and limited scalability, restricts its applications. The luminal dye CFSE presented with no detectable aggregates in suspension, supporting the idea that intravesicular labeling can avoid the aggregation artifacts characteristic of membrane dyes such as PKH26, underscoring the advantage of luminal dyes. Nevertheless, CFSE’s dependence on luminal esterases limits its broad applicability. While this enzymatic dependency can be beneficial for functional assays, it may limit the dye’s effectiveness for more general EV profiling. The inherent variation in esterase levels between EV populations likely accounts for the distinct CFSE-negative subset we resolved *via* EV-FCM and introduces variability that constrains its utility as an EV label. This heterogeneity aligns with previous reports describing weak or absent CFSE-labeling in EVs derived from cell types with low luminal esterase content, highlighting cell- and EV-type–specific limitations of this dye rather than a technical one^13^. In addition, we identified that CFSE labeling efficiency markedly differed between NKEVs (isolated *via* TFF-SEC) and 293EVs (isolated *via* UC). While EV-intrinsic esterase content likely contributes to this difference, the disparate backgrounds suggest that the isolation method itself may contribute to CFSE-labeling proficiency. UC co-sediments soluble proteins alongside EVs, whereas TFF-SEC preferentially removes soluble contaminants. Since soluble esterases present in EV preparations can activate CFSE non-specifically, a phenomenon documented in protein-enriched controls, UC may exacerbate this artifact compared to TFF-SEC. In contrast, CytoLight’s esterase-independent activation mechanism ensures consistent labeling across sample types and isolation methods, thereby improving signal-to-noise ratios and enabling more reliable quantification analyses across diverse EV sources^37^. CytoLight also preserves EV integrity and analytical accuracy, positioning it as a scalable, stable and cost-effective fluorophore suitable for quantitative and tracking applications both *in-vitro* and *in-vivo*^42^.

The second part of the study pursued evaluating CytoLight’s performance in functional studies where strong accurate fluorescence is essential. We identified 10µM as the saturation point that yields maximal vesicle loading and signal specificity. However, these concentrations yield high background signals attributable to unbound dye rather than to EV-associated fluorescence. Removal of this free dye is a necessary step for applications requiring the use of large quantities of labeled EVs, such as *in-vitro* EV uptake and *in-vivo* tracking. A brief UF step using regenerated cellulose membranes effectively removed ∼98% of free dye while preserving vesicle integrity and restoring signal specificity. Subsequent lysis of UF-cleaned samples with detergent validated that the remaining fluorescent events originated exclusively from intact vesicles and not from free dye in buffer or from co-isolated protein.

Building on our robust labeling and cleanup protocol, we evaluated CytoLight-labeled EV uptake. In flow cytometry-based uptake assays, cells exposed to CytoLight-labeled NKEVs displayed substantially higher fluorescence than cells treated with CytoLight-labeled PBS or with CytoLight-labeled, protein-matched EV-depleted serum, demonstrating that the major contribution to the cellular signal originates from EV-bound dye rather than from residual unbound dye or dye–protein complexes. This design explicitly addresses concerns previously raised by Takov *et al*., highlighting that protein and lipoprotein contaminants can confound uptake readouts when lipophilic dyes are used without stringent controls^43^. To our knowledge, our data represent the first application of such protein-matched controls to luminal dyes. In contrast to CytoLight, the additional luminal dye we tested, CFSE, did not generate EV-dependent uptake signals under our conditions, as CFSE-labeled NKEVs produced fluorescence indistinguishable from CFSE-labeled PBS despite UF or UC cleanup, demonstrating poor free-dye removal. Consequently, residual free CFSE molecules may be internalized by target cells and activated by intracellular esterases, becoming a significant source of non-specific cellular staining. This matches reports describing high background for CFSE in uptake assays^21,31^. In comparison, our UF-based approach for CytoLight cleanup overcomes these limitations and yields direct, rapid, reproducible, esterase-independent fluorescence that is immediately verifiable post-cleanup and suitable for downstream applications.

CytoLight-labeled EV uptake was consistent with an endocytosis-mediated pattern. Pre-treatment with the dynamin inhibitor Dynasore reduced CytoLight signal in both C6 and K562 cells, indicating endocytosis-mediated EV uptake confirming *bona fide* vesicle internalization. In alignment, confocal microscopy revealed a punctate intracellular pattern that colocalized with the endosomal marker TSG101, rather than a diffuse membranal distribution, further supporting endocytic trafficking of intact vesicles rather than simple dye adsorption under the conditions tested^44–46^. Within the constraints of our models, CytoLight provides a functionally interpretable uptake signal consistent with established endocytic uptake mechanisms. Determining whether the dye affects EV cargo functionality remains a critical next step. A potential approach would be to evaluate the preservation of NKEV-mediated cytotoxicity against K562 cells, as shown by Samara *et al*.^47^.

For *in-vivo* applications, CytoLight’s far-red emission spectrum, which minimizes tissue autofluorescence, proved advantageous^48^. Accordingly, mice injected intravenously with CytoLight-labeled EVs exhibited significantly higher fluorescence accumulation in the liver, colon, kidney, blood, lung and spleen, compared to controls receiving labeled PBS. This organ distribution is consistent with prior reports that intravenously administered EVs accumulate predominantly in liver and spleen and variably in lung and kidney depending on EV source and dose^49,50^. The differential signal between labeled EVs and labeled PBS indicates that CytoLight enables detection of EV-driven biodistribution rather than merely tracking free dye. While these *in-vivo* results are limited to one EV type, one model and two time points they demonstrate the practical utility of CytoLight for comparative biodistribution studied when appropriate controls are included.

Overall, CytoLight emerges as a superior alternative to current fluorescent dyes, combining high specificity, stability, minimal artifacts and ease of use with broad analytical compatibility. Its adherence to current best-practice recommendations and its ability to generate clean, quantifiable signals without extensive purification make it particularly suitable for both *in-vitro* and *in-vivo* EV studies. Together, these attributes position CytoLight as a reliable and broadly applicable tool for advancing standardized, aggregation-free EV characterization and tracking.

## Acknowledgements

We would like to express our appreciation to Rhenium, for providing access to their CytoFLEX nano flow cytometer used in the EV-FCM experiments, and extend our special thanks to Shlomit Rak Yahalom for her valuable guidance.

## Data Availability Statement

The data that support the findings of this study are available from the corresponding author upon reasonable request.

