## Supplementary figures for "CytoLight: A Rapid and Versatile Fluorescent-Based Labeling Method for Extracellular Vesicle Characterization and Tracking"

### Slide 1
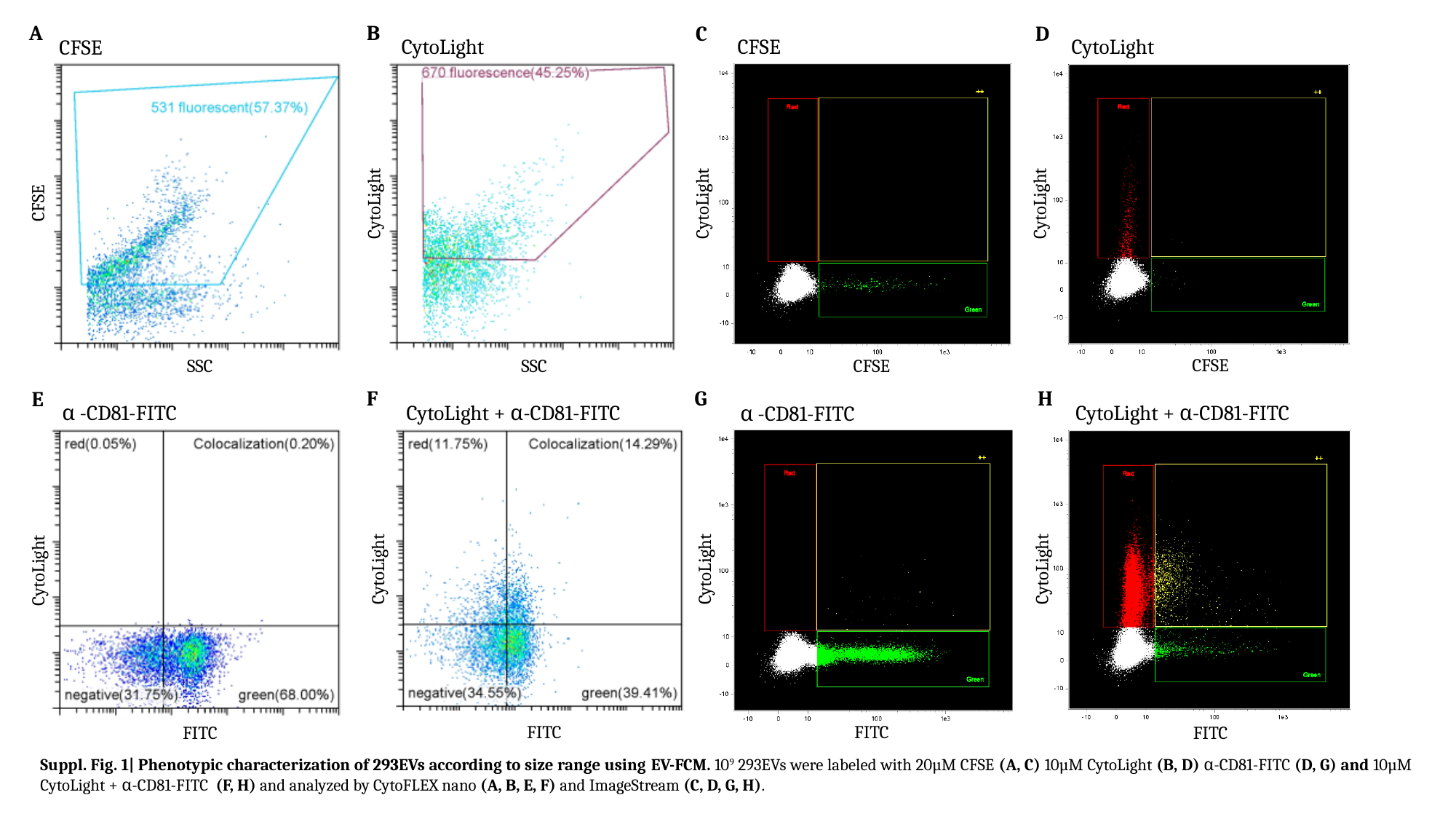

B
CytoLight
CytoLight
SSC
A
CFSE
CFSE
SSC
C
CFSE
CytoLight
CFSE
D
CytoLight
CytoLight
CFSE
G
α -CD81-FITC
CytoLight
FITC
H
CytoLight + α-CD81-FITC
CytoLight
FITC
F
CytoLight + α-CD81-FITC
CytoLight
FITC
E
α -CD81-FITC
CytoLight
FITC
Suppl. Fig. 1| Phenotypic characterization of 293EVs according to size range using EV-FCM. 10^9^ 293EVs were labeled with 20µM CFSE (A, C) 10µM CytoLight (B, D) α-CD81-FITC (D, G) and 10µM CytoLight + α-CD81-FITC (F, H) and analyzed by CytoFLEX nano (A, B, E, F) and ImageStream (C, D, G, H).

### Slide 2
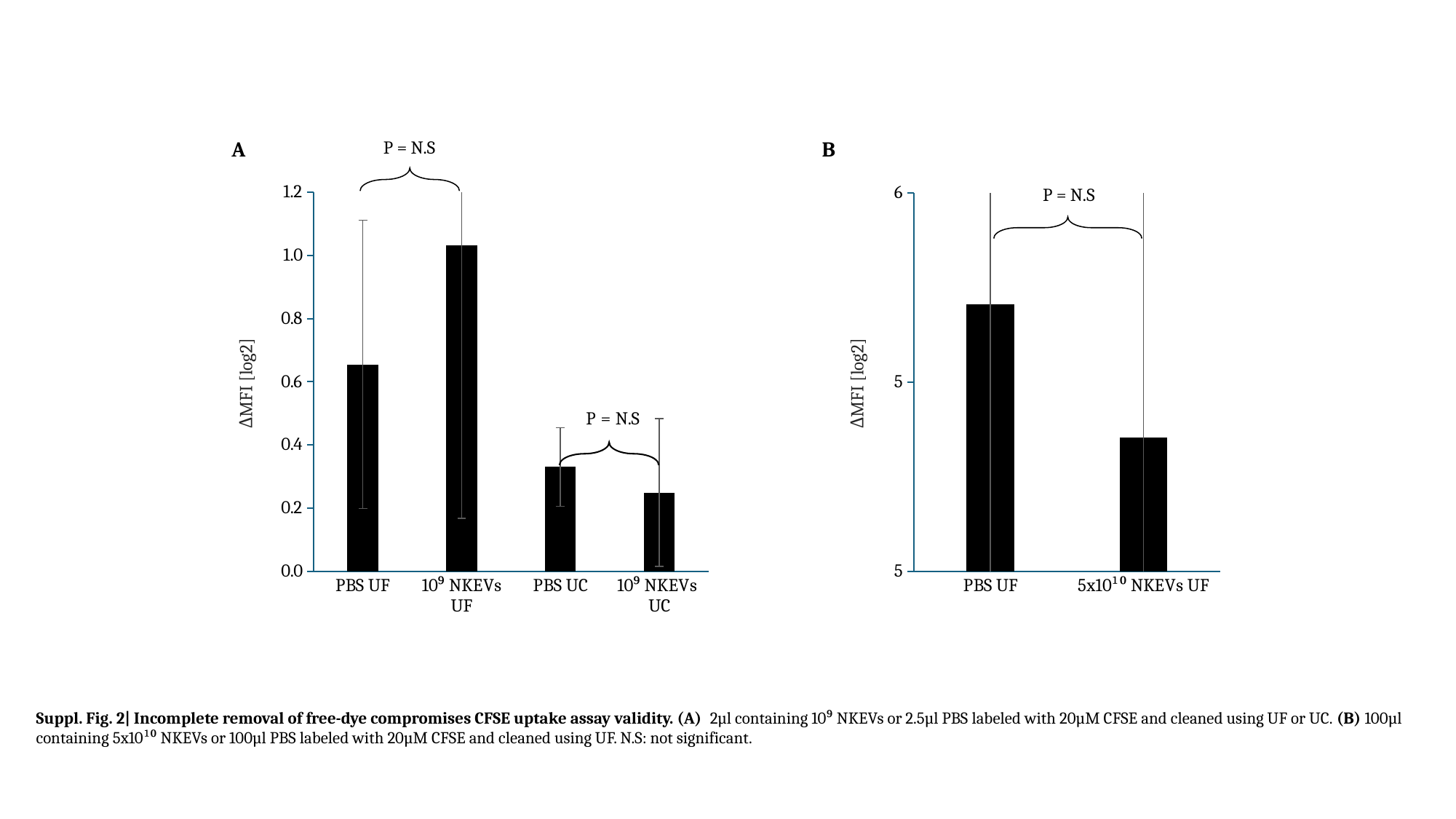

#### Chart
| Category | Log2fc over US |
|---|---|
| PBS UF | 0.654876159 |
| 10⁹ NKEVs UF | 1.032678356 |
| PBS UC | 0.3300161309948444 |
| 10⁹ NKEVs UC | 0.249139 |P = N.S
A
ΔMFI [log2]
#### Chart
| Category | Log2fc over US |
|---|---|
| PBS UF | 5.20564341556318 |
| 5x10¹⁰ NKEVs UF | 4.85294502722882 |B
P = N.S
ΔMFI [log2]
Suppl. Fig. 2| Incomplete removal of free-dye compromises CFSE uptake assay validity. (A) 2µl containing 10⁹ NKEVs or 2.5µl PBS labeled with 20µM CFSE and cleaned using UF or UC. (B) 100µl containing 5x10¹⁰ NKEVs or 100µl PBS labeled with 20µM CFSE and cleaned using UF. N.S: not significant.
